# Age-associated transcriptional programs in tumor-associated macrophages are associated with tumor progression in HPV-negative oral cancer

**DOI:** 10.64898/2026.10.02.756061

**Authors:** Hemant Gujar, Sarah Hanache, Sadhna Aggarwal

**Affiliations:** Columbia University Medical Center, Columbia University; Department of Leukemia, The University of Texas, MD Anderson Cancer Center; Department of Radiation Oncology, The University of Texas, MD Anderson Cancer Center

## Abstract

Aging and tobacco exposure are major determinants of risk and outcome in HPV-negative oral cavity squamous cell carcinoma (OSCC), yet the cellular programs linking host factors to tumor immunity remain poorly defined. We reprocessed and reanalyzed previously published single-cell RNA-seq data from 19 HPV-negative OSCC tumors and we identified a robust age-and smoking-associated expansion of SPP1^high^ tumor-associated macrophages (TAMs), accompanied by a shift toward a myeloid-dominant and lymphocyte-reduced tumor microenvironment. SPP1^high^ TAMs exhibit a distinct transcriptional state enriched for extracellular-matrix remodeling and metabolic programs and reduced antigen-presentation pathways. They also express inflammatory chemokines such as CXCL2 and CXCL3 and down-regulate the cytotoxic mediator TNFSF10, suggesting a broadly protumor TAM phenotype. Ligand-receptor modeling suggests that SPP1^high^ TAMs act as key communication hubs, with dominant SPP1-CD44 signaling to T-cell subsets. Transcription-factor analysis further implicates a POU2F1/POU2F2/SP1 module in maintaining this state. In the TCGA HNSCC cohort, the association between SPP1 expression and overall survival did not reach statistical significance after adjustment for age and tumor purity. These findings demonstrate that aging and tobacco exposure converge to expand an SPP1^high^ TAM population that remodels the extracellular matrix and suppresses antitumor immunity, highlighting the SPP1-CD44 axis and its upstream regulators as potential therapeutic targets in high-risk OSCC.

## Introduction

Oral cancer, that includes cancer of the lips, tongue, and oral cavity, represents about 3% of all cancers in the United States, with an estimated 54,540 new cases and 10k deaths every year (1) As per the GLOBOCAN report, globally, oral cancer is one of the most prevalent cancers, with approximately 3.7 lakh new cases and 1.7 lakh deaths reported each year (2). Globally, it is more prevalent in countries like India, Thailand, and Bangladesh, where the use of tobacco, betel nut and alcohol are major risk factors, contributing to oral cancer rates that are >3 times in Western countries. One of the key challenges in oral cancer treatment is its advanced stage diagnosis when the tumor has spread to surrounding tissues and lymph nodes, making surgery and radiation less effective. Despite advances in research and technology that has developed new targeted therapies and immunotherapies, the prognosis remains poor for many patients, underscoring the need for earlier detection and more personalized treatment strategies (3, 4). Additionally, the genetic complexity and heterogeneity of oral cancers present significant obstacles to effective treatment, requiring ongoing research to better understand tumor biology and develop more effective therapies (5, 6). Recent studies emphasize the need for early detection markers and more immunotherapy options as promising advances in improving outcomes (7). Among oral squamous cell carcinomas, HPV-negative tumors account for the majority of oral cavity cancers and are characterized by poorer clinical outcomes, greater genomic complexity, and strong associations with tobacco exposure and aging.

SPP1 (Secreted Phosphoprotein 1), also known as osteopontin, is a glycoprotein that plays a significant role in cell adhesion, migration, and survival (8). It is involved in several biological processes, including immune responses, inflammation, and tissue remodeling, making it particularly relevant in cancer progression (9–11). In tumors, SPP1 expression is upregulated, contributing to tumor metastasis, angiogenesis, and poor prognosis due to its promotion of tumor cell invasion (12, 13). It interacts with integrin and CD44 receptors on various cells, influencing both tumor and immune cells within the tumor microenvironment (9). Consequently, SPP1 is under investigation as a potential diagnostic, prognostic, and therapeutic marker for cancer immunotherapy (14–16).

The tumor-associated macrophages (TAMs) play a crucial role by imparting immune-interaction in tumor immuno-microenvironment (TIME) that contributes towards tumor’s fate. SPP1^high^ macrophages represent an important TAM cluster in several types of cancers (15), heart diseases (17), tissue fibrosis (18, 19), nonalcoholic steatohepatitis (20) etc. Higher SPP1-macropahges have been observed in several tumor types like colorectal cancer, gastric, breast, liver, NSCLC and PDAC cancers (21–24). SPP1 expression and secretion regulate macrophage biology as it promotes macrophage polarization toward either M1 or M2 like TAMs (25, 26). It induces the hypoxia-inducible factor 1α (HIF-1α) expression leading to M1 polarization in colorectal cancer (27, 28). In nonalcoholic fatty liver disease also, these macrophages promote M1 polarization by stimulating of the JAK1/STAT1/HMGB1 axis that activates the secretion of HMGB1 (29). However, in HCC the SPP1^high^ macrophages induced the polarization of macrophages to M2-TAM (30) by regulating CD44 and PTGER4 signaling axis. As per a recent study, macrophages expressing both SPP1 and CXCL9 upregulated M0, M1, and M2 markers in cancers, but SPP1High macrophages showed higher M2 marker profile (31). SPP1^high^ macrophages express macrophage polarization associated factors such as C1QC, MRC1, STAT1, and PPARG, reflecting that SPP1 play a significant role in TAM polarization (27). In ESCC, abundance of SPP1^high^ SIRPα+ macrophages have been found to be positively associated with patient’s responsiveness to PD-1/PD-L1 inhibitors by inducing PD1+CD8+ T cells, indicating a potential role for these macrophages in shaping the immune landscape and treatment outcomes (32). Functionally, SPP1 induces TAMs to profile as Inflammatory-TAMs and/or proangiogenic TAMs that either induces inflammatory cytokine production thereby recruiting anti-tumor immune cells or promotes tumor EMT and angiogenesis respectively (21, 33). A recent study of HNSCC clinical samples (n=5) by scRNA-seq data analysis revealed that SPP1^high^ TAMs correlated with poor prognosis in these patients by promoting cell proliferation and migration, secreted higher levels TNF-α and IL-1β and induced NF-kappa B signaling axis. They also observed that these cells promoted the expression of SPP1 in both tumor cells and other neighboring macrophages as a feedback loop via unknown mechanisms (16).

In aging tissues, the accumulation of SPP1^high^ macrophages has been observed, especially in fibrotic and degenerative conditions This study showed that SPP1 regulated macrophage recruitment during the senescence of fibro-adipogenic progenitors (34). In chronic kidney disease (CKD) and heart failure, SPP1^high^ macrophages expand and correlate with fibrosis, indicating their role in disease progression (33). Interestingly, SPP1^high^ TAMs are more prevalent in older individuals, possibly due to the cumulative effects of aging-related inflammation and tissue remodeling (35).

Despite increasing recognition of SPP1^high^ macrophages in cancer, how aging and tobacco exposure influence the emergence and functional state of these macrophages in HPV-negative oral squamous cell carcinoma (OSCC) remains poorly understood. To address this question, we reanalyzed published single-cell RNA-seq data from HPV-negative OSCC to determine how aging and tobacco exposure shape macrophage heterogeneity, with particular emphasis on SPP1^High^ tumor-associated macrophages and their transcriptional programs and intercellular communication.

## Results

### Immune and stromal cell composition of the HPV-negative OSCC microenvironment

Single-cell profiling of HNSCC revealed that the complexity of the tumor microenvironment plays a central role in disease progression(36) yet its cellular and molecular complexity in relation to aging and environmental risk factors remain poorly characterized. To characterize the tumor microenvironment in HPV-negative OSCC, we reprocessed and reanalyzed single-cell RNA-seq data from 19 HPV-negative OSCC tumors (19 patients) originally generated by Kürten et al. (37) (NCBI SRA: SRP301444).

Sample-level clinical annotations (sex, smoking status, alcohol use, inflammation category, and TNM stage) were obtained from the accompanying dataset metadata and the primary publication and used for downstream stratified analyses (Table 2). The cohort initially comprised seven young and twelve old patients (Supplementary Figure S2A). For immune-focused analyses, samples with fewer than 200 immune cells were excluded to ensure robustness, resulting in six young and six old samples with sufficient immune-cell representation included in downstream comparisons. Age groups were defined using a prespecified cutoff of 65 years old. This cutoff was selected to capture clinically relevant age-associated immune remodeling. ScRNASeq data was analyzed using Cell Ranger (38). Samples were corrected for background RNA contamination using (CellBender: (39)). Droplets with more than one cell (doublets) were removed using Scrublet (40) and Doublet detection (41). Cells with mitochondrial RNA content greater than 5% and number of genes less than 500 were removed using Seurat (42, 43) for further down-stream analysis. The average of cells per samples were 2059. The median mitochondrial content in samples ranged from 0.5 % to 2.65 %; median ribosomal RNA content in samples ranged from 13.4 % to 28.5 %; median number of genes identified in sample ranged from 1286 to 3312; and the median number of RNA molecules in samples ranged from 2389 to 10533. UMAP dimensionality reduction and clustering using Seurat, identified 15 distinct clusters (Supplementary Figure S1). Cells were first stratified based on PTPRC expression into immune (PTPRC+) and non-immune (PTPRC-) cells. Re-clustering of non-immune cells revealed eight cell types. Basal cell (KRT14, S100A2), endothelial cell 1 (CCL21, SEMA3A), fibroblast (LUM, COL1A1), endothelial cell 2 (PLVAP, COL15A1), smooth muscle cell (RGS5), plasma cell (IGKC, IGHG1), cancer cell (WFDC2, KRT19), and myocyte (ACTA1, SLN). Re-clustering of immune cells resolved 12 clusters including T cell 00 (LTB, BCL11B), T cell 01 (CXCL13)-which are T follicular helper or exhaustion-like CD4+ T-cell states-, Macrophage (S100A9, S100A8), T cell 03 (P2RY8), T cell 04 (GNLY, KLRF1), plasma cell (BANK1, CD79A), monocyte (JARID2, ATP2B1), plasmacytoid dendritic cells (DC) cluster 1 (JCHAIN, TCF4), heat-shock-responsive myeloid cluster (HSPA1A, DNAJB1), myeloid DC (TXN, FSCN1), basophil (TPSAB1, TPSB2), and plasmacytoid DC cluster 2 (CPVL). (Figure 1a-d). Marker genes used for cell-type annotation and macrophage subclustering are provided in Supplementary Tables S1 and S3.

**Figure 1.**
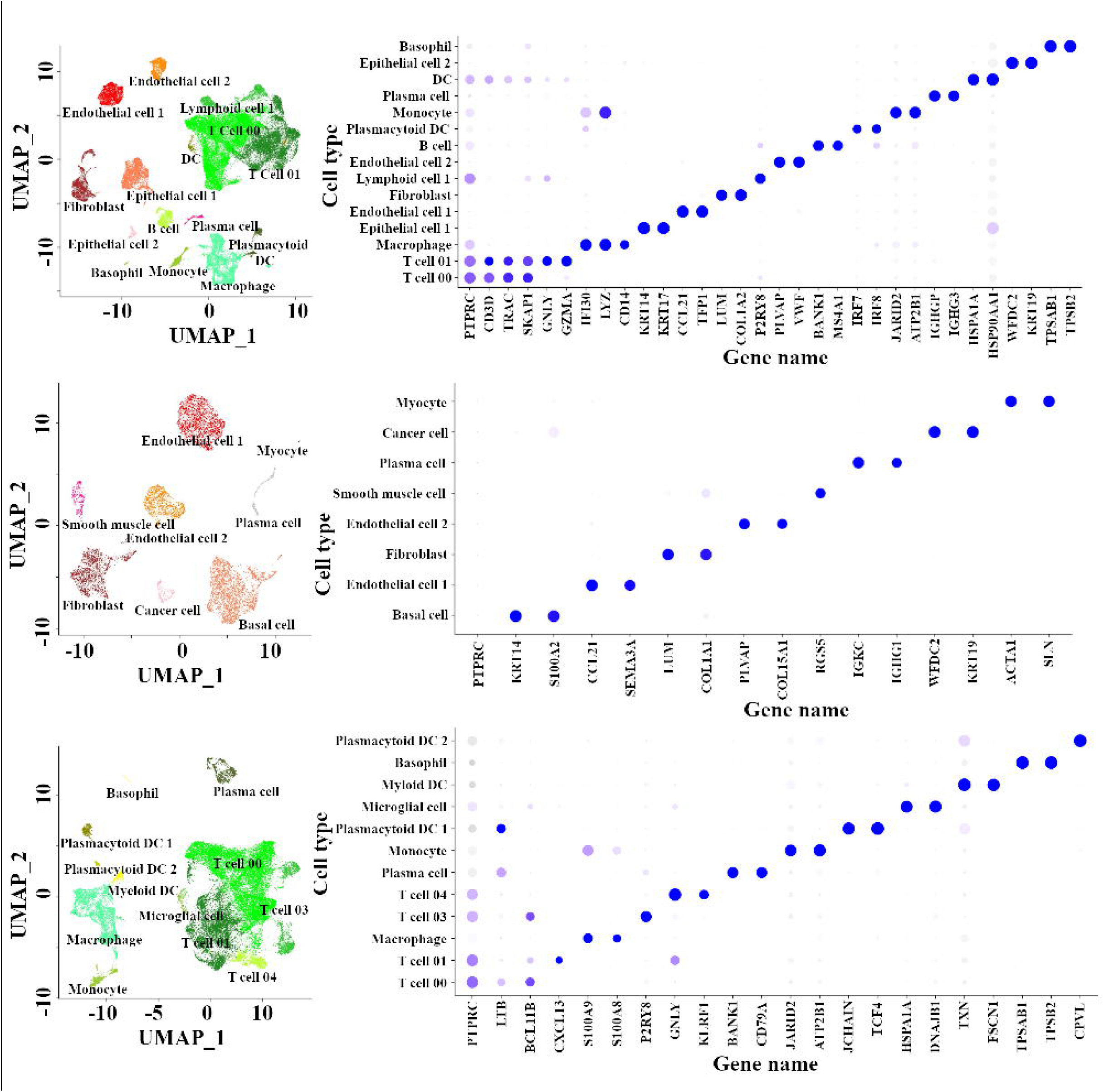
Cellular diversity and marker gene expression in HNSCC (a) UMAP of re-clustered CD45-negative cells. (b) Dot plot showing marker gene expression in CD45-negative cell clusters. (c) UMAP of re-clustered CD45-positive cells. (d) Dotplot showing marker gene expression in CD45-positive immune clusters.

**Table 1.** Multivariable Cox proportional hazards model evaluating the prognostic effect of *SPP1* expression adjusted for age and tumor purity.

| Variable | Coefficient | Hazard Ratio (HR) | SE (coef) | 95% CI Lower | 95% CI Upper | Z-value | P-value | Significance |
| --- | --- | --- | --- | --- | --- | --- | --- | --- |
| *SPP1* | 0.051 | 1.052 | 0.029 | 0.995 | 1.113 | 1.786 | 0.074 |  |
| Age | 0.023 | 1.023 | 0.006 | 1.01 | 1.036 | 3.534 | <0.001 | *** |
| Purity | 0.15 | 1.162 | 0.318 | 0.624 | 2.166 | 0.474 | 0.636 |  |

**Table 2.**
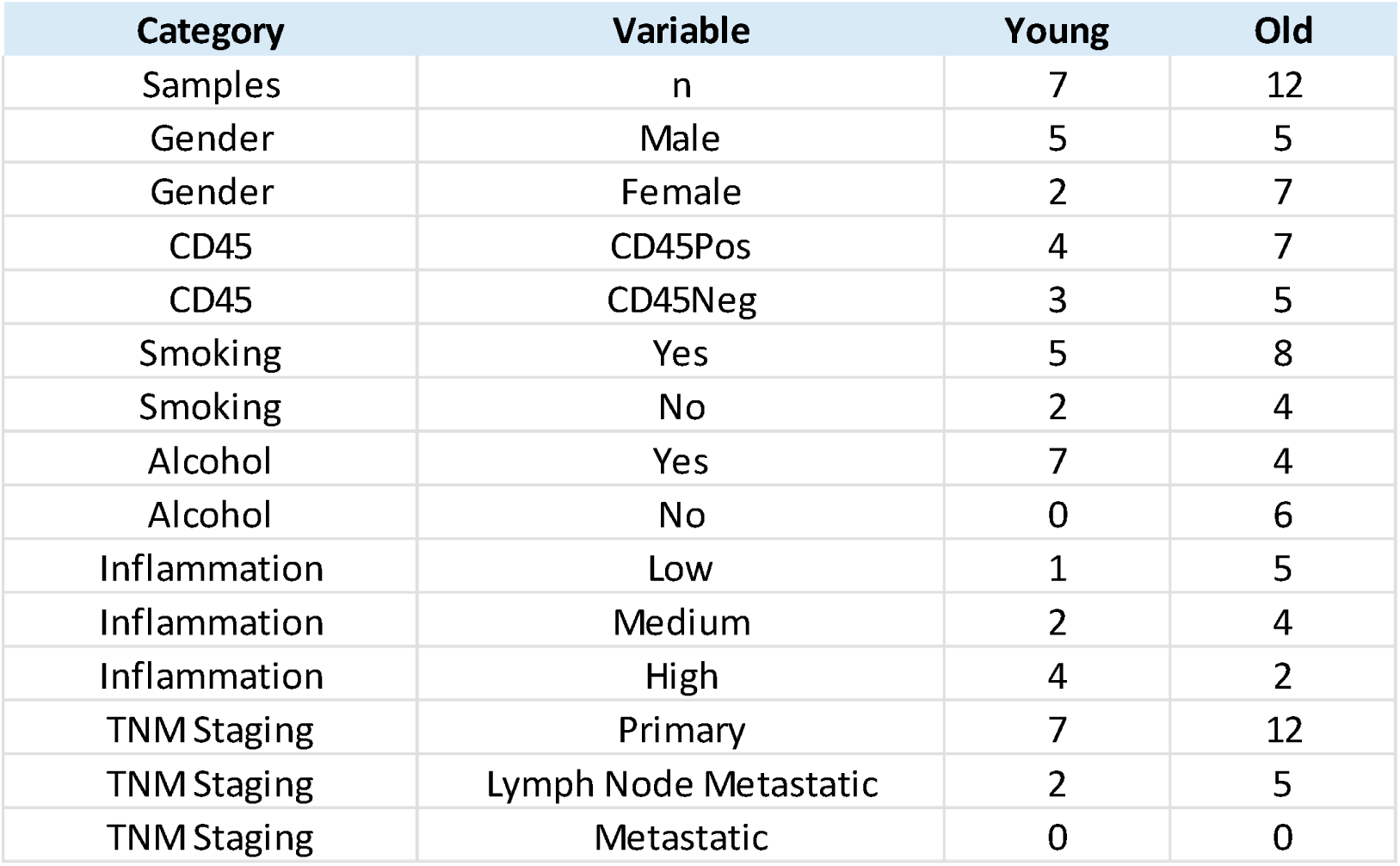
Clinical and demographic characteristics of the original HPV-negative OSCC cohort.

### Aging Shifts the Landscape of Tumor Microenvironment Towards Myeloid-Dominance

In solid cancers, aging was shown to promote myeloid skewing and impair immune functions, accentuating tumorigenesis, and increasing mortality. (44) (45) In HNSCC, tumor-associated macrophages are a major source of PD-L1 and other immune checkpoint ligands(37). However, the changes in molecular and cellular composition of TME in aging is not well characterized. To assess age-associated immune remodeling in HNSCC, we analyzed immune cells from six young (age<65) and six old patients (age >65) with a total of 14604 cells in both the groups. Samples GSM5017062, GSM5017041, GSM5017023, GSM5017044, GSM5017050, GSM5017025 from old group contained a limited number of immune cells and were discarded for this analysis. Older patients exhibited increased frequencies of CXC13+ T cells (consistent with T follicular helper or exhaustion-like CD4+ T-cell states T cell 01), macrophage, basophil, and plasmacytoid DC2. Whereas the number of P2RY8+ cytotoxic T cells (T cell 03), monocyte, plasma cell, plasmacytoid DC1, and myeloid DC were enriched in younger as compared to older population. T cell 00 and T cell 04 remained relatively stable across age groups. T cell 03 and monocyte cells were observed mostly in younger population only (Figure 2). CXCL13+ T cells (consistent with T follicular helper or exhaustion-like CD4+ T-cell states) are often associated with dysfunctional states (36) while P2RY8+ T cells (cytotoxic CD8+ T-cell populations) exhibit activated cytotoxic features, suggesting that HNSCC in older patients may be less surveilled by cytotoxic T cells and more prone to immune evasion (46). These findings suggest that aging shifts the immune landscape of HNSCC favoring dysfunctional T cell subsets potentially linked to immune dysfunction. Seurat FindMarkers function with default settings and a cutoff of min.pct = 0.40; logfc threshold = 0.25 was used to identify differential gene expression in all immune cell clusters in old and young samples. At a cut-off of p-adjusted value 0.05, 31 genes were recorded in cluster 0, 188 genes in cluster 1, 182 genes in cluster 2, 11 genes in cluster 3, 170 genes in cluster 4, 37 genes in cluster 5, 30 genes in cluster 6, 52 genes in cluster 7, 165 genes in cluster 8, 26 genes in cluster 9, 6 genes in cluster 10, and 4 genes in cluster 11 were observed (Supplementary Table S1). Re-clustering of TAMs identified six transcriptionally distinct subsets defined by dominant marker genes including IDO1, C1Q, S100A12, SPP1, CCL3L1 and TCF7L2 (Fig 3a., 3b.). Notably, the SPP1^high^, CCL3L1^high^ and C1Q^high^ subsets were disproportionately enriched in older patients, with 85%, 67% and 79% of respective cells originating from aged individuals (Fig 3c., 3d.).

**Figure 2.**
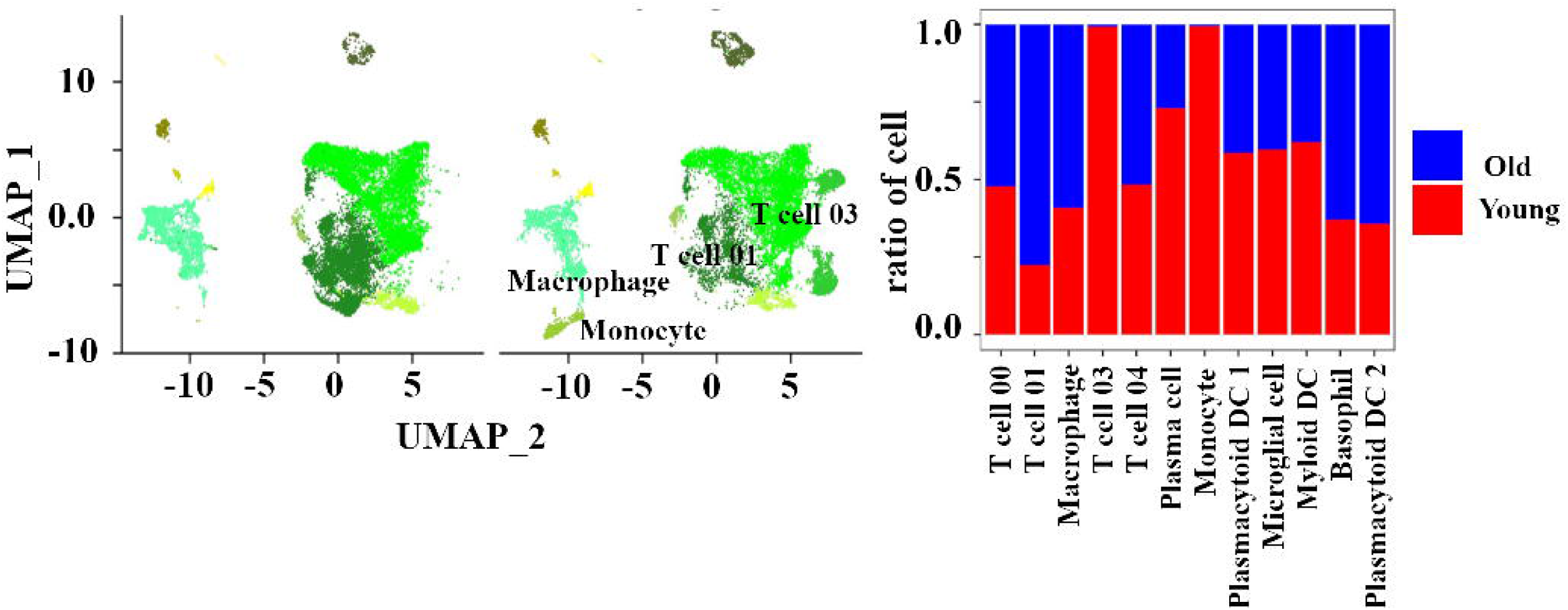
Age-associated shifts in immune cell populations (a) UMAP of immune cells from young and old patient samples. (b) Proportional distribution of immune cell types by age group.

**Figure 3.**
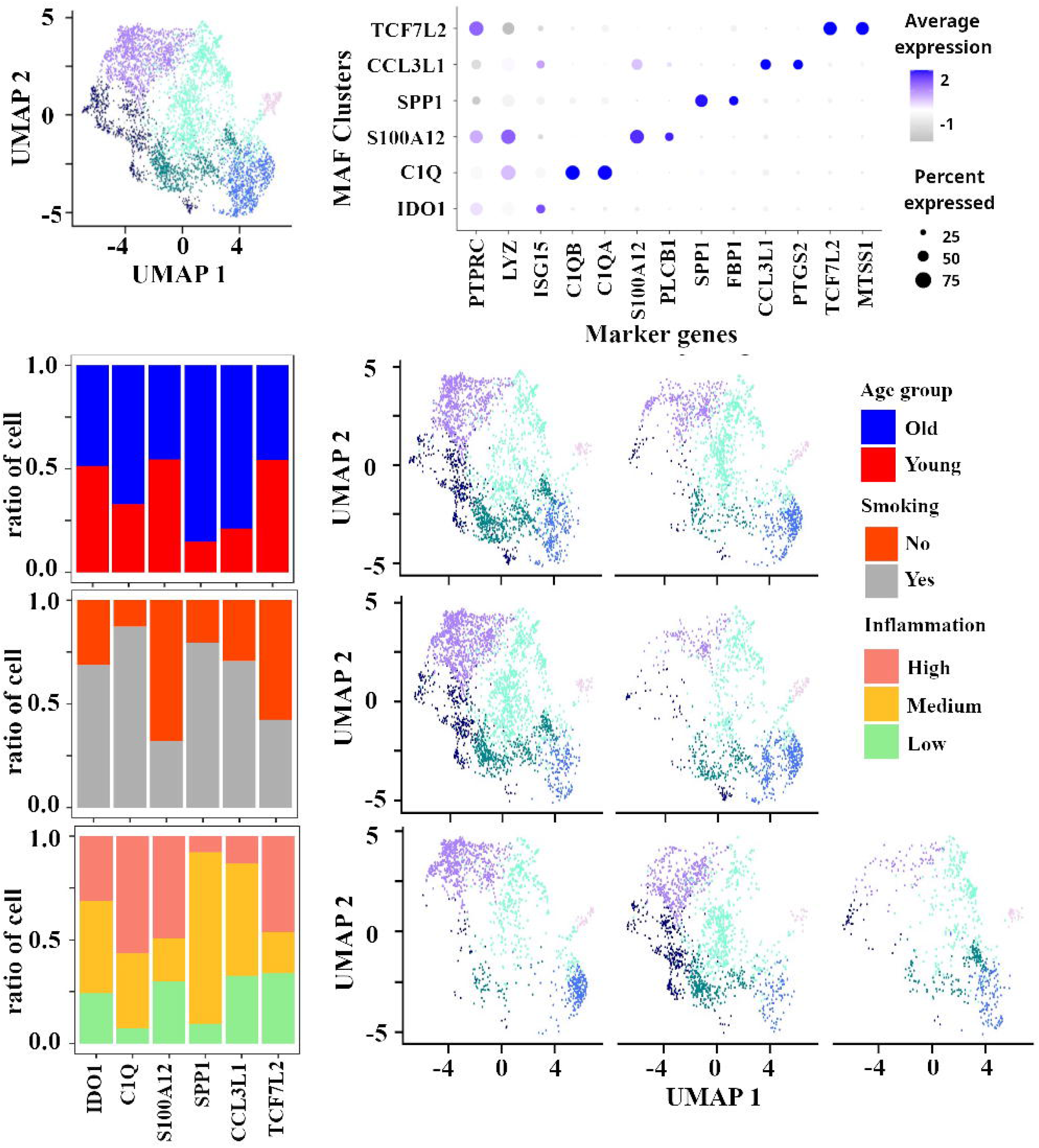
Tumor-associated macrophage subsets and their distribution (a) UMAP of TAM subclusters identified by reclustering macrophages. (b) Dot plot showing marker genes defining each TAM subset. (c) Proportion of TAM subsets in young and old samples. (d) UMAP showing TAM subset distribution by age. (e) Proportion of TAM subsets in smokers and non-smokers. (f) UMAP showing TAM subset distribution by smoking status. (g) Distribution of TAM subsets across low, intermediate, and high inflammation groups. (h) UMAP showing TAM subset distribution by inflammation level.

Tobacco exposure further influenced macrophage subtype distributions: SPP1^high^, CCL3L1^high^ and IDO1^high^ TAMs are increased in smokers while S100A12^high^ and TCF7L2+ subsets were prevalent in non-smokers (3e, 3f). Importantly, SPP1^high^ TAMs were absent from highly inflamed tumors, being confined to low and moderate inflammation settings-Inflammation was assigned based on histopathological annotation provided in Kürten et al. 2021-(3g, 3h). In the TCGA HNSCC cohort (n = 492), the association between SPP1 expression and overall survival did not reach statistical significance after adjustment for age and tumor purity (HR = 1.052, 95% CI: 0.995-1.113, P = 0.074). This data suggests that SPP1^high^ TAMs may actively contribute to the establishment of “immune cold” TME characterized by reduced T-cell infiltration. This supports previous reports showing that SPP1-expressing myeloid cells contribute to T cell dysfunction and immune evasion in lung adenocarcinoma and renal cell carcinoma through SPP1 and CD44 mediated signaling and extracellular matrix remodeling. Moreover, SPP1^high^ TAMs represent a core immunoregulatory module enriched in invasive, therapy resistant and low immune-infiltrated tumors(31, 47). Together with our findings, these studies reinforce a model in which SPP1^high^ TAMs might exhibit an immune-poor and tumor-permissive microenvironment in the context of HNSCC.

### SPP1 TAMs Exhibit Pro-Tumorigenic and Immunosuppressive Gene Programs

Differential genes expressed in SPP1 ^high^ TAM vs other TAMs were recorded using Seurat FindMarkers function with default settings. Differential gene expression between SPP1high and SPP1low TAMs identified 762 significant genes (adjusted p < 0.05), including 607 upregulated and 155 downregulated genes (Supplementary Table S4). Among the most significantly upregulated genes in SPP1 TAMs were FN1, FBP1, MIF, MARCO. These genes, identified in Supplementary Table S4, are consistently associated with extracellular matrix remodeling (FN1), metabolic adaptation (FBP1), and immunomodulatory signaling (MIF, MARCO)(31, 48–51), supporting a coordinated pro-tumorigenic macrophage program.(Fig. 4a, b). These features point to a macrophage population specialized for immune silencing rather than inflammatory response. Samples were stratified in two groups, according to quantile-based thresholds, high SPP1 samples (SPP1^high^) with normalized SPP1 RNA > 2 (Top 25%), number of cells > 200 / ratio of # of cells with high SPP1 expression >= 0.25 and low SPP1 samples (SPP1^low^) with normalized SPP1 RNA close to 0 (bottom 25%) and number of cells > 200 / ratio of # of cells with low SPP1 expression >= 0.75. All SPP1^high^ samples were predominantly derived from older patients (GSM5017022 (HN01), GSM5017025 (HN02), GSM5017040 (HN08), GSM5017043 (HN09)). Samples in SPP1^low^ group contained both the old (GSM5017027 (HN03), GSM5017061 (HN15)) and young (GSM5017031 (HN05), GSM5017046 (HN10)) Age group samples (Figure 4).

**Figure 4.**
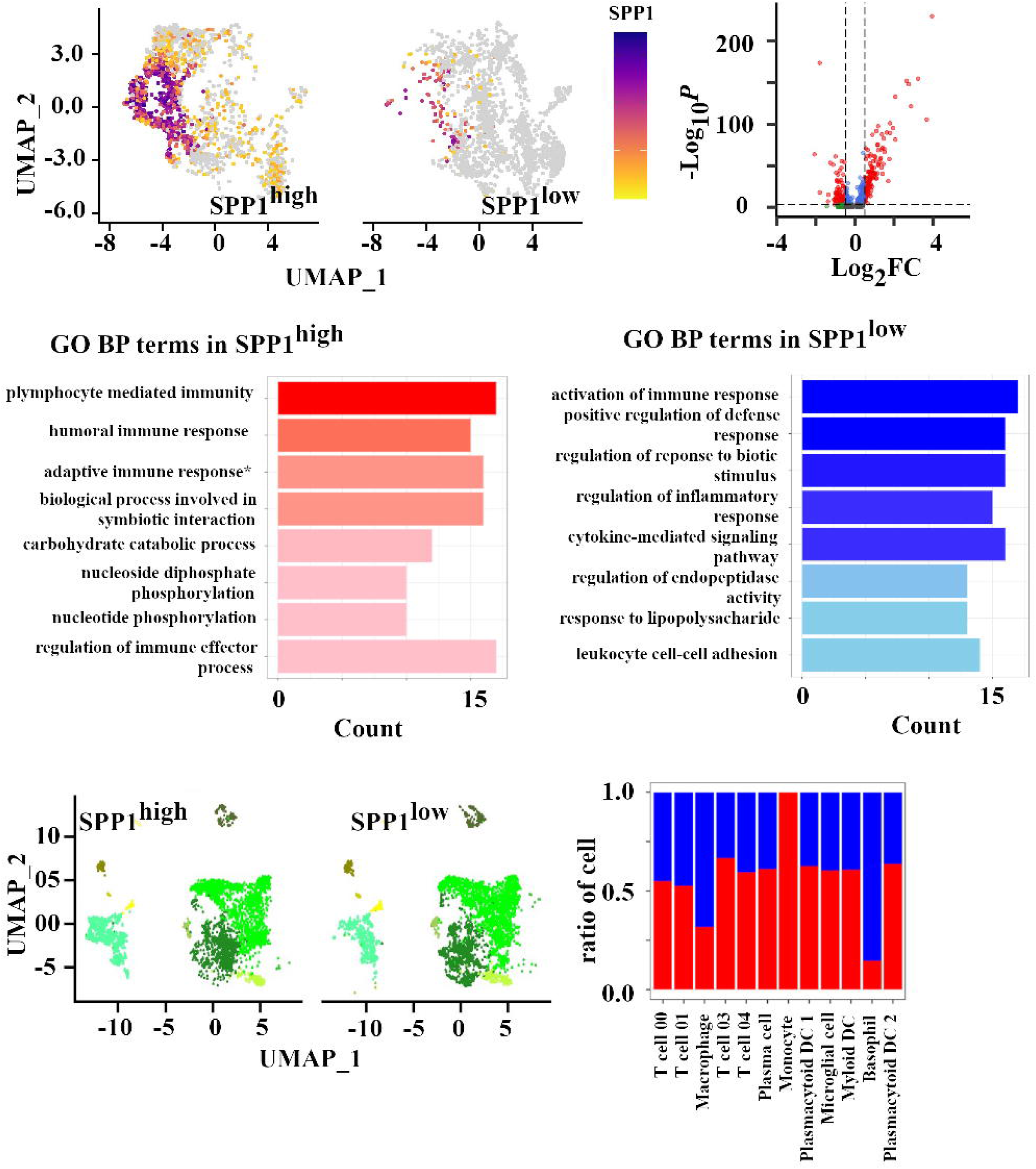
SPP1^high^ versus SPP1^low^ tumor-associated macrophages (TAMs) (a) Feature plot showing SPP1 expression in TAMs. (b) Volcano plot of differentially expressed genes between SPP1^high^ and SPP1^low^ TAMs. (c) Gene ontology enrichment of differentially expressed genes. (d) UMAP showing immune cell distribution in SPP1^high^ and SPP1^low^ samples. (e) Proportions of immune cell types in SPP1^high^ versus SPP1^low^ groups. (f) Heatmap of M1/M2-associated macrophage gene scores in SPP1high and SPP1low TAMs.

Genes up-regulated in SPP1^high^ macrophage including FBP1, APOC1, MMP12, GPNMB and APOE have previously been associated with tumor progression, macrophage polarization, and extracellular matrix remodeling in multiple cancer types. (50)

Gene ontology analysis showed that genes elevated in SPP1^high^ TAMs were associated with B cell signaling, humoral immunity, lymphocyte regulation, and metabolic processes including nucleotide and carbohydrate metabolism (Fig. 4c). In contrast, downregulated genes were involved in inflammatory signaling, cytokine activity, and leukocyte adhesion, suggesting a shift away from classical immune effector functions. We next assessed immune composition in tumors enriched for SPP1^high^ TAMs. Samples with high SPP1 expression showed increased macrophage and basophil content and reduced frequencies of T cells and dendritic cells (Fig. 4d, e). These findings indicate that SPP1^high^ TAMs are associated with a myeloid-dominant immune landscape and diminished lymphocyte infiltration. Since GO terms related to immune process were enriched in the differentially expressed genes, ratio of different immune population was recorded. Comparison of number of immune cells in SPP1^high^ and SPP1^low^ samples with 978 cells randomly selected from every sample to ensure equal cell representation across groups showed that macrophage and basophil population were higher in SPP1^high^ group whereas, T cell population and other remaining cell populations were marginally higher in SPP1^low^ group (Figure 4e).

### Aging and smoking associate with transcriptional states that converge on SPP1^high^ TAM phenotypes

To determine whether environmental or host factors influence the emergence of SPP1^high^ TAMs, we compared gene expression profiles in TAMs from older versus younger individuals and smokers versus non-smokers. A total of 377 genes were differentially expressed with age and 641 with smoking exposure (Fig. 5a, b) (supplementary table 5). Of these, 67% and 45%, respectively, overlapped with the SPP1 TAM signature (Fig. 5c, d), including canonical SPP1-associated ECM and inflammatory mediators (VCAN, CXCL2, IL1B) indicating a shared transcriptional program (31). These data suggest that both intrinsic aging and extrinsic exposures promote a macrophage phenotype associated with immunosuppression and poor T cell recruitment. Gene ontology enrichment analysis of overlapping genes revealed significant enrichment in extracellular matrix organization, cytokine signaling, and inflammatory response pathways (Supplementary Table S5). Notably, classical TAM markers such as SPP1, VCAN, and CXCL2 were enriched in older individuals whereas smokers exhibited increased expression of C1QC, C1QA, C1QB, ISG15, CXCL2, CCL3, and IL1B. These findings indicate that aging and smoking are associated with transcriptional programs that are consistent with suppressive or remodeling TAM phenotypes. Importantly, SPP1^high^ TAMs were found exclusively in tumors with low or intermediate levels of inflammation and were absent in highly inflamed samples (Fig. 5e), linking their presence to a tumor microenvironment with diminished immune infiltration. Together, these results suggest that both biological aging and tobacco exposure favor the accumulation of SPP1^high^ TAMs within immunologically “cold” tumors, potentially contributing to impaired anti-tumor immunity.

**Figure 5.**
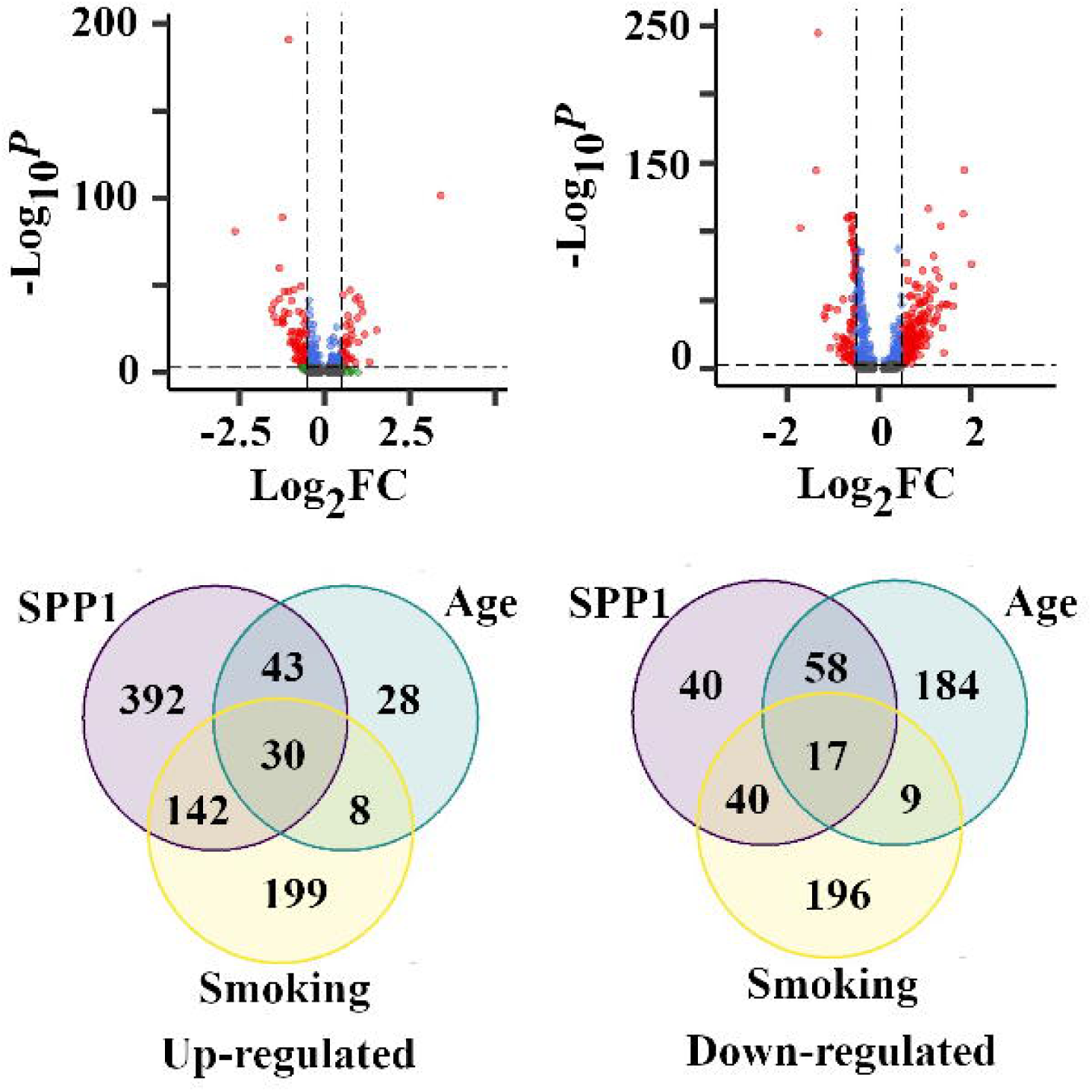
Shared transcriptional programs in aging, smoking, and SPP1 status (a) Volcano plot of gene expression differences in TAMs from older versus younger patients. (b) Volcano plot of gene expression differences in TAMs from smokers versus non-smokers. (c) Venn diagram showing overlap of upregulated genes in SPP1^high^ TAMs, aging, and smoking. (d) Venn diagram showing overlap of downregulated genes in SPP1^high^ TAMs, aging, and smoking. (e) showing overlapping differentially expressed genes shared between aging-, smoking-, and SPP1-associated macrophage programs

### Immunosuppressive transcriptional regulatory program and cytokine profile in SPP1^high^ TAMs

To identify transcriptional regulators driving SPP1^high^ TAM state, we compared transcription factor activity in SPP1^high^ vs SPP1^low^ TAMs using decoupleR framework. The top 25 differentially active TFs distinguishing the two groups were visualized in a heatmap. (Fig. 6a). Among these *SP1*, *POU2F1*, and *POU2F2* showed markedly increased activity and expression in the SPP1^high^ macrophage, with increased inferred activity scores (*POU2F1* = 0.38, *POU2F2* = 0.34, *SP1* = 0.31; Supplementary Table S5a (Fig 6b), indicating a coordinated transcriptional program underlying the SPP1^high^ phenotype.” These TFs are known to regulate genes involved in extracellular matrix remodeling, immune regulation, and macrophage activation and to bind the SPP1 promoter region and regulate SPP1 transcription in other cancer and cellular systems. (51, 52) These findings suggest that the POU-SP1 transcriptional module may drive the emergence of immunosuppressive SPP1^high^ TAMs in the aging HPV negative oral cancer microenvironment.

**Figure 6.**
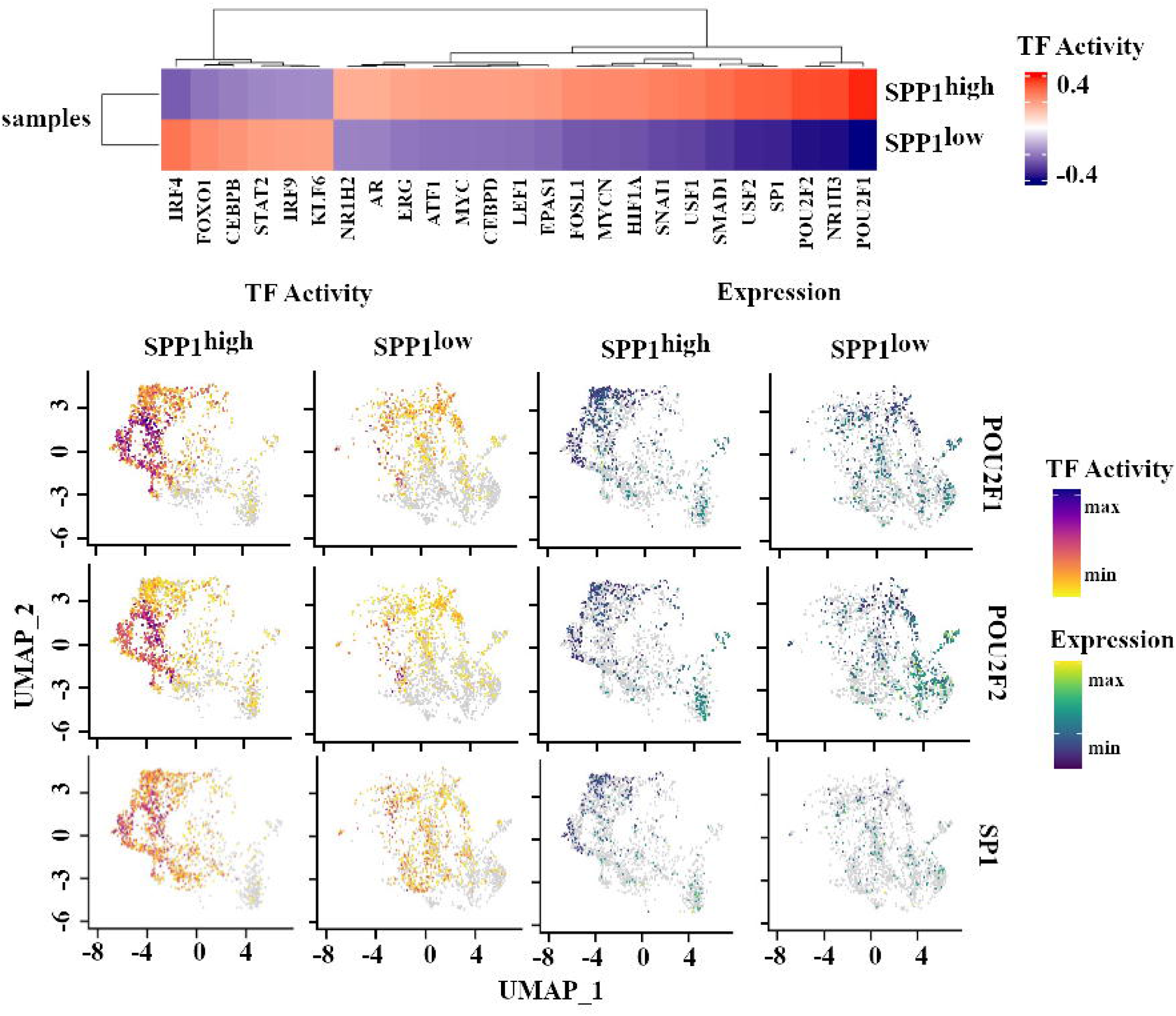
Transcription factor activity in SPP1^high^ TAMs (a) Heatmap of the top 25 transcription factors with differential activity between SPP1^high^ and SPP1^low^ TAMs. (b) UMAP plots showing inferred transcription factor activity and gene expression for POU2F1, POU2F2, and SP1.

### SPP1^high^ TAMs Orchestrate Immunosuppressive Signaling and Tumor Progression Through Underlying Transcriptional Programs

To elucidate the functional phenotype of SPP1^high^ TAMs, we conducted a differential gene expression between SPP1^high^ and SPP1^low^ TAMs. This analysis revealed up-regulation of genes associated with extracellular matrix, remodeling and immune modulation, including SPP1, FN1, MIF, FBP1, and MARCO. (Supplementary Table S4). These genes have been implicated in promoting tumor invasiveness and creating an immunosuppressive microenvironment in various cancers. Given the up-regulation of genes linked to immune suppression and extracellular matrix remodeling in SPP1^high^ TAMs, we hypothesized that this subset may actively modulate the tumor microenvironment through altered cytokine expression. 19 cytokine-related genes (Supplementary Table S4) were observed to be differentially expressed between SPP1^high^ TAMs and other TAM subsets, with CXCL8 and CXCR4 upregulated, while the majority, including the cytotoxic mediator TNFSF10 (TRAIL), were downregulated, consistent with an immunosuppressive macrophage phenotype. TNFSF10 has been shown to exert anticancer effects and suppress tumor development in murine models of skin cutaneous melanoma in mice. (53) In transgenic mice, targeted expression of TNFSF10 protects against chemically induced skin carcinogenesis (54). Moreover, low TNFSF10 expression in human melanoma correlates with aggressive tumor features and reduced immune infiltration, suggesting that loss of TNFSF10-mediated signaling may facilitate immune escape (53). Additionally, SPP1^high^ TAMs exhibited increased expression of integrins ITGA5 and ITGAM, along with FN1, which are associated with enhanced cell adhesion and migration, further supporting their role in facilitating tumor progression(55). Collectively, these findings underscore the multifaceted role of SPP1^high^ TAMs in promoting tumor growth and suppressing anti-tumor immunity, highlighting them as potential targets for therapeutic intervention. Difference in transcription factor activity was recorded in TAMs in the SPP1^high^ group vs SPP1^low^ group using R package decoupleR (56) using default parameters. Transcription factor activity and expression of selected genes POU2F1, POU2F2, and SP1 were plotted in FeaturePlot_scCustom (Figure 6b).

### Predicted SPP1-CD44 signaling links SPP1^high^TAMs to T-cell subsets

To assess the functional impact of SPP1^high^ TAMs on immune crosstalk, we used CellChat to infer ligand-receptor signaling between TAM subsets and T-cell clusters using default parameters (Figure 7). We focused on the ‘Secreted Signaling’ category in the CellChat human database. (39.6% of curated interactions) to capture diffusible signaling between cell types. Network visualization of the number of inferred interactions (Figure 7a) and interaction weights (Figure 7b) revealed extensive macrophage-T cell communication, with prominent connectivity involving the SPP1^high^ (mSPP1) and C1Q^high^ (mC1Q) TAM populations across multiple T-cell subsets. Decomposition of these interactions into individual ligand-receptor contributions demonstrated that the SPP1-CD44 axis was the dominant driver of macrophage-T cell communication (Figure 7c). The next most influential interactions were mediated by MIF signaling through CD7 co-receptor complexes, including MIF (CD74+CD44) and MIF (CD74+CXCR4), indicating that macrophage-T cell crosstalk is reinforced by multiple immunoregulatory pathways. In addition, several lower-amplitude interactions reflected broader SPP1-centered signaling via integrin receptors (e.g., SPP1(ITGA4+ITGB1), SPP1(ITGAV+ITGB1), SPP1(ITGA5+ITGB1), and SPP1(ITGA9+ITGB1)), alongside chemokine axes such as CCL5-CCR1, CCL3-CCR1, and CXCL13-CXCR3. Together, these results nominate an SPP1-centric communication program dominated by SPP1-CD44 and supported by MIF-CD74 pathways as a principal mode of TAM-to-T cell signaling, consistent with SPP1^high^ TAMs functioning as key communication hubs within the OSCC microenvironment.

**Figure 7.**
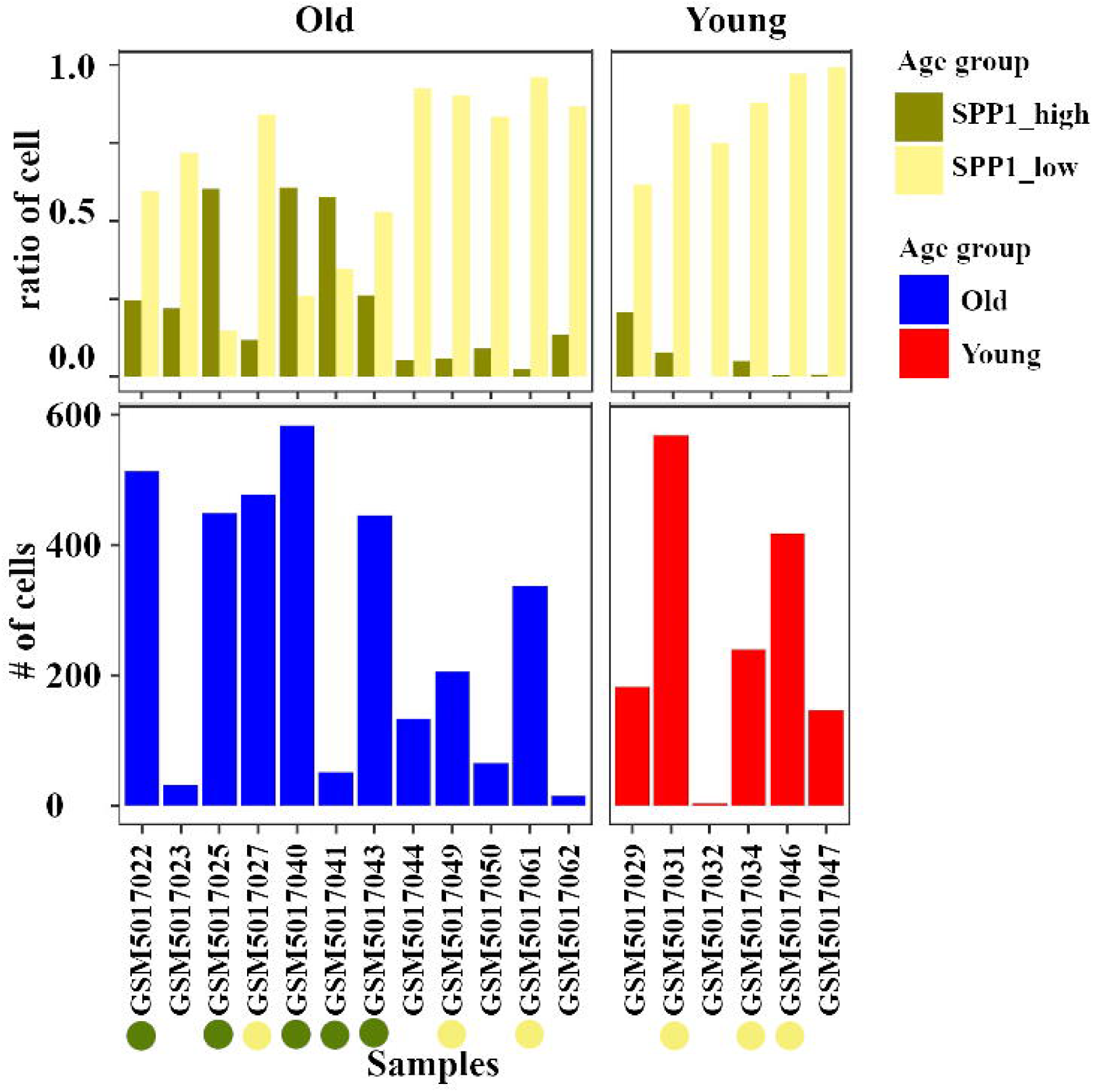
Ligand-receptor communication between TAMs and T-cell subsets (a) Total number of ligand-receptor interactions between TAMs and T-cell subsets. (b) Weighted interaction strength between macrophage and T-cell subsets. (c) Contribution of major ligand-receptor pairs, including the SPP1-CD44 axis, to macrophage-T-cell communication.

At the global level, communication patterns revealed that these TAMs contributed disproportionately to outgoing signaling modules, including those related to matrix remodeling and cytokine signaling (Figure 8). These findings position SPP1^high^ TAMs as central hubs in the tumor microenvironment, with potential relevance to immune evasion and therapy resistance.

**Figure 8.**
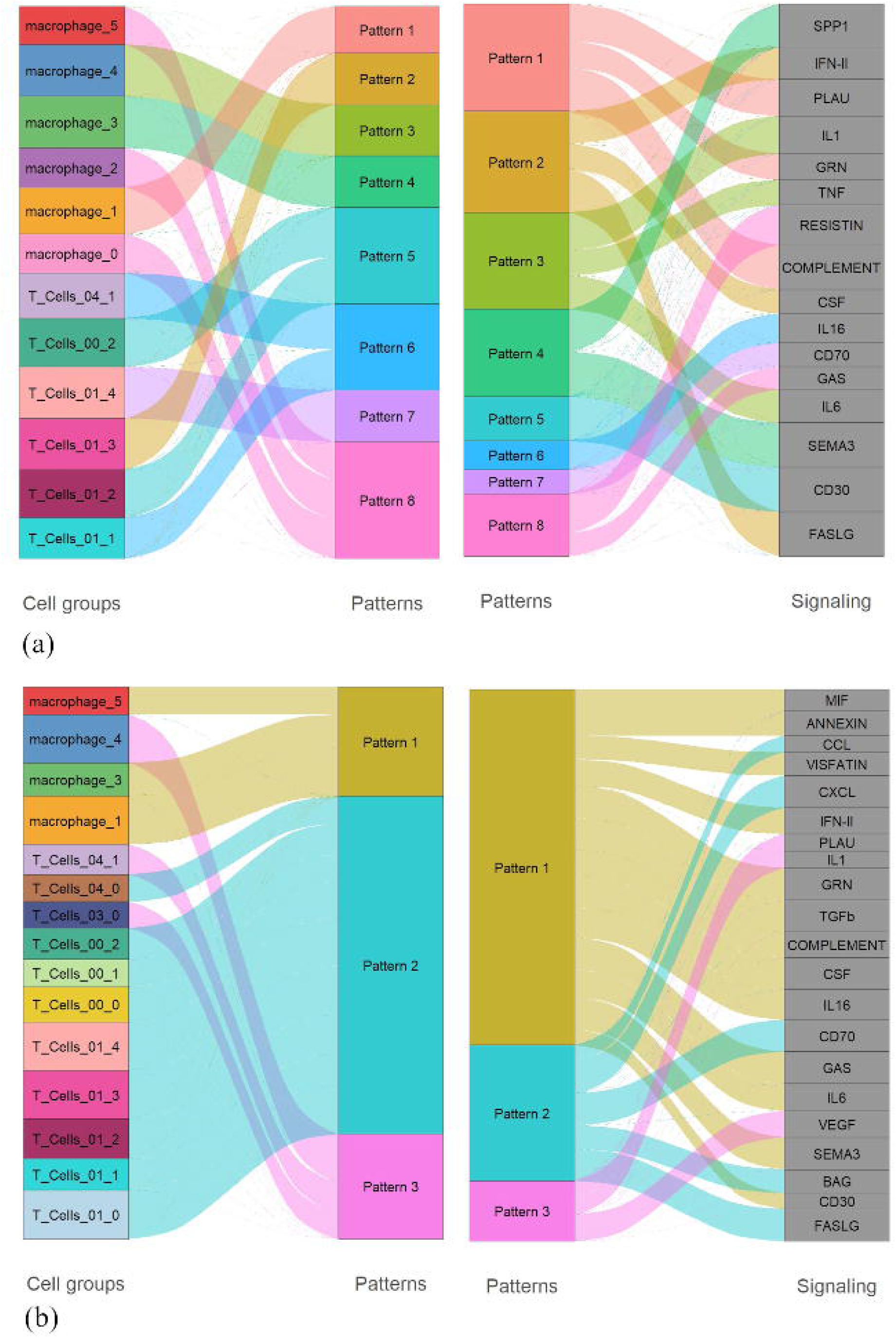
Global signaling patterns between TAMs and T cells (a) River plot showing incoming signaling pathways grouped by communication pattern. (b) River plot showing outgoing signaling pathways contributed by macrophage and T-cell subsets.

## Discussion

In this study, our single-cell RNA-seq analysis of HPV-negative oral HNSCC revealed that aging and smoking associate with a myeloid-dominant tumor microenvironment enriched in SPP1^high^ tumor-associated macrophages (TAMs).

SPP1 has recently been shown to be an active driver of tumor aggressiveness. In pancreatic ductal adenocarcinoma, genetic ablation of SPP1 reprogrammed cancer cells toward an epithelial phenotype and suppressed metastasis, establishing SPP1 as essential for maintaining mesenchymal cell fate. (57) SPP1^high^ macrophages have emerged as a conserved immuno-regulatory subset across cancer types, characterized by extracellular-matrix remodeling, inflammatory cytokine secretion, and poor clinical outcome. (47) Such macrophages represent a distinct lineage repeatedly observed in fibrosis, heart failure, and malignancy, underscoring their role as chronic inflammation sensors. (58) Moreover, recent studies reporting that SPP1 orchestrates pro-tumor immune microenvironment across cancers by engaging CD44-dependent signaling to remodel the extracellular matrix and suppress cytotoxic T-cell activity. (47, 59)

SPP1^high^ TAMs were transcriptionally distinct, with enrichment of extracellular-matrix organization and lipid-metabolism programs, including FN1, MIF, and FBP1, together with reduced antigen-presentation and cytokine-response modules. Comparable SPP1-associated macrophage programs have been reported in hepatocellular and colorectal cancers, including interactions with fibroblasts that promote immune exclusion. Our findings extend these observations to HPV-negative oral HNSCC. (51, 60, 61)

Beyond ECM and metabolic genes, we observed strong upregulation of CXCL2 and CXCL3 in SPP1^high^ TAMs. CXCL2 is associated with poor prognosis in ovarian cancer (62) and promotes tumor progression and metastasis in glioblastoma (63). CXCL3 has been implicated in hepatocellular cancer stem-cell maintenance (64), uterine cervical cancer progression (65), and pancreatic cancer aggressiveness (49). Their coordinated induction suggests that SPP1^high^ TAMs adopt a broadly pro-tumor inflammatory program with known roles across multiple malignancies.

We further demonstrate, using the analyzed scRNA-seq cohort that SPP1^high^ TAMs act as bidirectional communication hubs with T-cell subsets, predominantly through the SPP1-CD44 axis. Comparable macrophage-T-cell immunosuppressive interactions have been reported in pancreatic and brain tumors.

Our transcription-factor activity analysis identified increased activity of POU2F1, POU2F2, and SP1 within the SPP1^high^ TAMs, supporting a mechanistic model in which a POU-SP1 regulatory module amplifies SPP1 expression and sustains an immunosuppressive macrophage phenotype. Consistent with this model, the human SPP1 promoter has been shown to contain functional binding sites for OCT-1 (POU2F1), OCT-2 (POU2F2), and SP1 that cooperatively regulate its transcription in tumor cells.(66)

Integrating our single-cell results with clinical metadata, we observed that all SPP1^high^ samples originated from older or smoking-exposed patients, whereas SPP1^low^ group contained both young and old individuals. It reinforces population-level and pan-cancer evidence linking SPP1 expression with aging, chronic inflammation, and immunosuppressive macrophage programs. (35)

Our dataset further reveals that SPP1^high^ TAMs have lower expression of TNFSF10 (TRAIL), consistent with the tumor-suppressive and immune-supportive roles of TRAIL reported in skin carcinogenesis and melanoma. In a transgenic mouse model, keratinocyte-restricted TRAIL over-expression protects against chemical skin carcinogenesis, indicating that TRAIL can mediate anti-tumor surveillance *in vivo*. (54) In human cutaneous melanoma, low TNFSF10 expression was associated with adverse clinicopathological features, reduced immune infiltration, and poorer benefits from anti-PD-1 or anti-CTLA-4. (53, 54) Together, these data support the idea that loss of TNFSF10 in SPP1^high^ TAMs reflects a shift away from cytotoxic, tumor-restraining programs toward more reparative or immunosuppressive macrophage states during ageing or exposure-driven inflammation.

Additionally, IL7R was upregulated in SPP1^high^ TAMs. While IL7R is best known for its role in lymphoid biology, gain-of-function IL7R mutations initiate ALL, highlighting the oncogenic potential of dysregulated IL7R signaling. Its induction here may indicate aberrant inflammatory activation or atypical macrophage-lymphoid crosstalk in aged/exposed tissues.

The nonsignificant survival association in TCGA limits conclusions about the prognostic value of SPP1. These integrative analyses are original to this work and corroborate pan-cancer observations linking SPP1^high^ macrophage modules with immune-cold, therapy-resistant states, as described in multiple SPP1 ^high^ TAM studies(16, 57). This study has several limitations. The analyses are based on a retrospective reanalysis of a publicly available single-cell RNA-seq dataset with a relatively modest cohort size after subgroup stratification. In addition, transcription factor activity and ligand-receptor interactions were computationally inferred and therefore require experimental validation. Finally, the cross-sectional study design identifies associations between aging, tobacco exposure, and macrophage states but does not establish causal relationships.

Collectively, our data establish that ageing and tobacco exposure converge to expand an SPP1^high^ TAM population that orchestrates ECM remodeling, represses antigen presentation, and dampens T-cell immunity. Targeting the SPP1-CD44 axis or its upstream transcriptional regulators (POU2F1/2, SP1) could re-program TAMs and restore immune surveillance in aged or high-risk oral cancer patients. These findings identify SPP1^high^ TAMs as a potential clinically relevant myeloid state that integrates environmental, age-related, and transcriptional drivers of immune suppression in oral cancer.

## Methods

### Study cohort and clinical annotation

This study is a secondary analysis of a previously published single-cell RNA-seq dataset comprising 19 HPV-negative oral cavity squamous cell carcinoma (OSCC) tumors from 19 patients originally reported by Kürten et al. (37). Raw sequencing data were downloaded from the NCBI Sequence Read Archive (SRA accession SRP301444). Clinical annotations including age, smoking history, alcohol use, inflammation category, and TNM staging were obtained from the associated metadata and are summarized in Table 2. HPV-negative status was defined in the primary study. For immune-focused analyses, samples containing fewer than 200 immune (CD45+) cells were excluded, resulting in six young and six old samples retained for downstream comparisons. Cluster-defining marker genes used for annotation are provided in Supplementary Tables S1 and S3.

### Single-cell RNA sequencing and preprocessing

Raw FASTQ files were processed with Cell Ranger v6.1.2 using default parameters and aligned to GRCh38. To reduce background RNA contamination, filtered feature-barcode matrices were processed using CellBender remove-background (39). Doublets were identified using Scrublet (40) and DoubletDetection (41) and removed prior to downstream processing.

Cells were retained if they contained >500 detected genes and <5% mitochondrial RNA using Seurat v4.3 (42). Across samples, the median mitochondrial content ranged from 0.5% to 2.65%, median ribosomal RNA content from 13.4% to 28.5%, median detected genes from 1286 to 3312, and median UMI counts from 2389 to 10,533. Each sample contained an average of 2059 high-quality cells. Clinical characteristics of the original HPV-negative OSCC cohort and quality-control metrics for the single-cell RNA-seq data are shown in Supplementary Figure S2.

### Normalization, integration, and dimensionality reduction

Filtered cells were normalized using NormalizeData() with a scale factor of 10,000 and log transformed. Gene-level variance was stabilized using FindVariableFeatures(), and samples were integrated using Seurat’s reciprocal PCA integration workflow to control for donor-level batch effects.

Principal component analysis (PCA) was performed on the integrated matrix, and UMAP was computed using the first 30 PCs. Cells were clustered using the Louvain algorithm with FindNeighbors() and FindClusters() at resolution 0.6 unless stated otherwise.

### Cell-type annotation

Major immune and stromal lineages were identified using canonical marker genes (e.g., CD3D for T cells, LYZ for myeloid cells, COL1A1 for fibroblasts, PECAM1 for endothelial cells). Manual refinement was guided by SingleR reference-based mappings and comparison to known immune signatures. Myeloid subclustering was performed by subsetting macrophages and re-clustering using the same PCA-UMAP-Louvain workflow.

### Differential expression and pathway enrichment

Differentially expressed genes (DEGs) between SPP1high and SPP1low TAMs were identified using Seurat’s FindMarkers() (Wilcoxon rank-sum test), with genes considered significant at adjusted p < 0.05 and |log2FC| > 0.5. Gene Ontology (GO) enrichment was performed on significantly expressed genes using clusterProfiler, highlighting pathways related to extracellular matrix remodeling, lipid metabolism, inflammatory signaling, and antigen presentation.

To compare immune composition between SPP1 groups, each sample was downsampled to a random set of 978 cells. Immune cell proportions were computed using Seurat cluster identities.

### Transcription factor activity analysis

Transcription factor (TF) regulatory activity was inferred using decoupleR (56) with DoRothEA regulons. TF activity scores were computed for each macrophage and compared between SPP1high and SPP1low TAMs. Expression and activity of TFs were visualized using FeaturePlot_scCustom.

### Cell-cell interaction analysis

Cell-cell interaction was inferred using CellChat v1.5, using the human ligand-receptor interactome. Interaction strength, outgoing and incoming signaling, and network centrality were computed across major cell lineages.

### TCGA HNSCC analysis

Bulk RNA-seq and clinical metadata for TCGA HNSCC tumors (n = 492) were downloaded from the Genomic Data Commons. TCGA analysis was restricted to HPV-negative HNSCC cases. SPP1 expression was correlated with CD8+ T-cell infiltration, showing a negative association (Supplementary Figure S2). Overall survival was analyzed using a multivariable Cox proportional hazards model including SPP1 expression, age, and tumor purity.

### Statistics

All statistical analyses were performed in R v4.2.0. Differential gene expression between cell populations was performed using the Wilcoxon rank-sum test implemented in Seurat. Differences in immune cell proportions between groups were evaluated using Fisher’s exact test, and nonparametric tests (Wilcoxon rank-sum or Kruskal-Wallis) were applied where appropriate. Multiple hypothesis testing was corrected using the Benjamini-Hochberg method. P values < 0.05 were considered statistically significant unless otherwise specified. Because cells derived from the same tumor are not statistically independent, clinical associations involving age, smoking status, and survival were interpreted at the patient level, whereas differential expression analyses were used to characterize transcriptional differences between cellular populations within the cohort.

Supplementary materials include: Figure S1 (cell-type marker genes), Table S1 (age-associated differential expression), Table S2 (TAM marker genes), Table S3 (SPP1^high^ vs SPP1^low^ macrophage DEGs), and Table S4 (transcription factor activity scores).

## Ethics Statement

This study involved secondary analysis of de-identified, publicly available human single-cell RNA-seq data. Ethical approval and informed consent for sample collection and sequencing were reported in the primary study (37). No additional institutional review board approval was required for this reanalysis.

---

**Figure S1.**
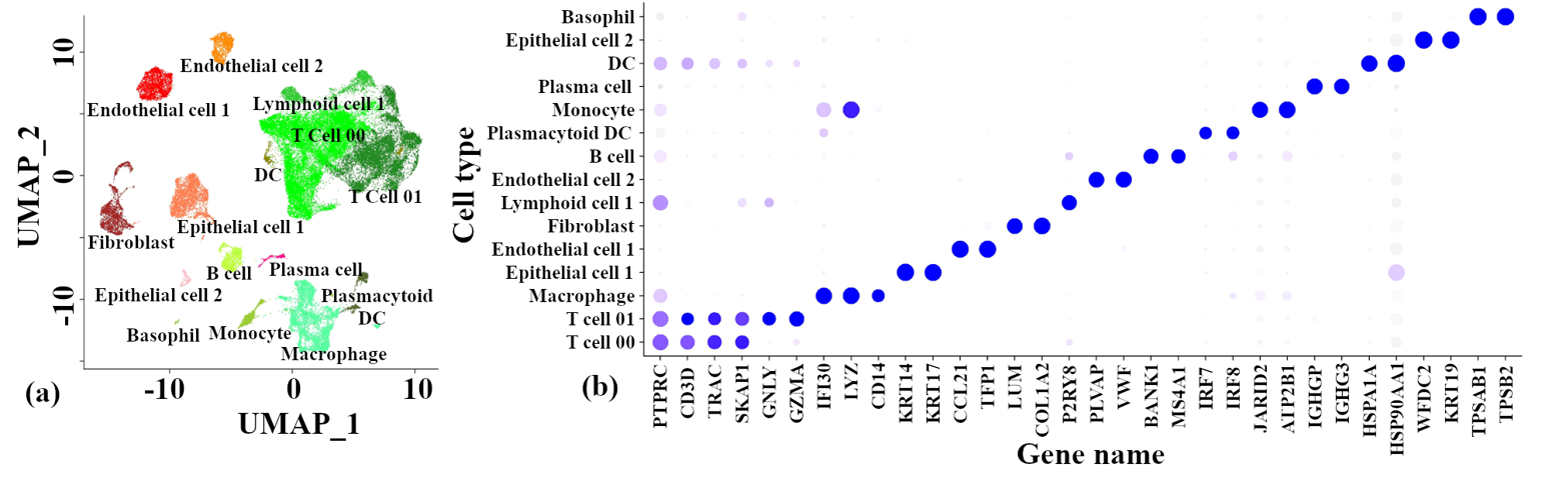
(a) UMAP plot showing major cell types identified by Seurat clustering of single-cell RNA sequencing data. (b) Dot plot showing the expression of selected marker genes across annotated cell types.

**Figure S2.**
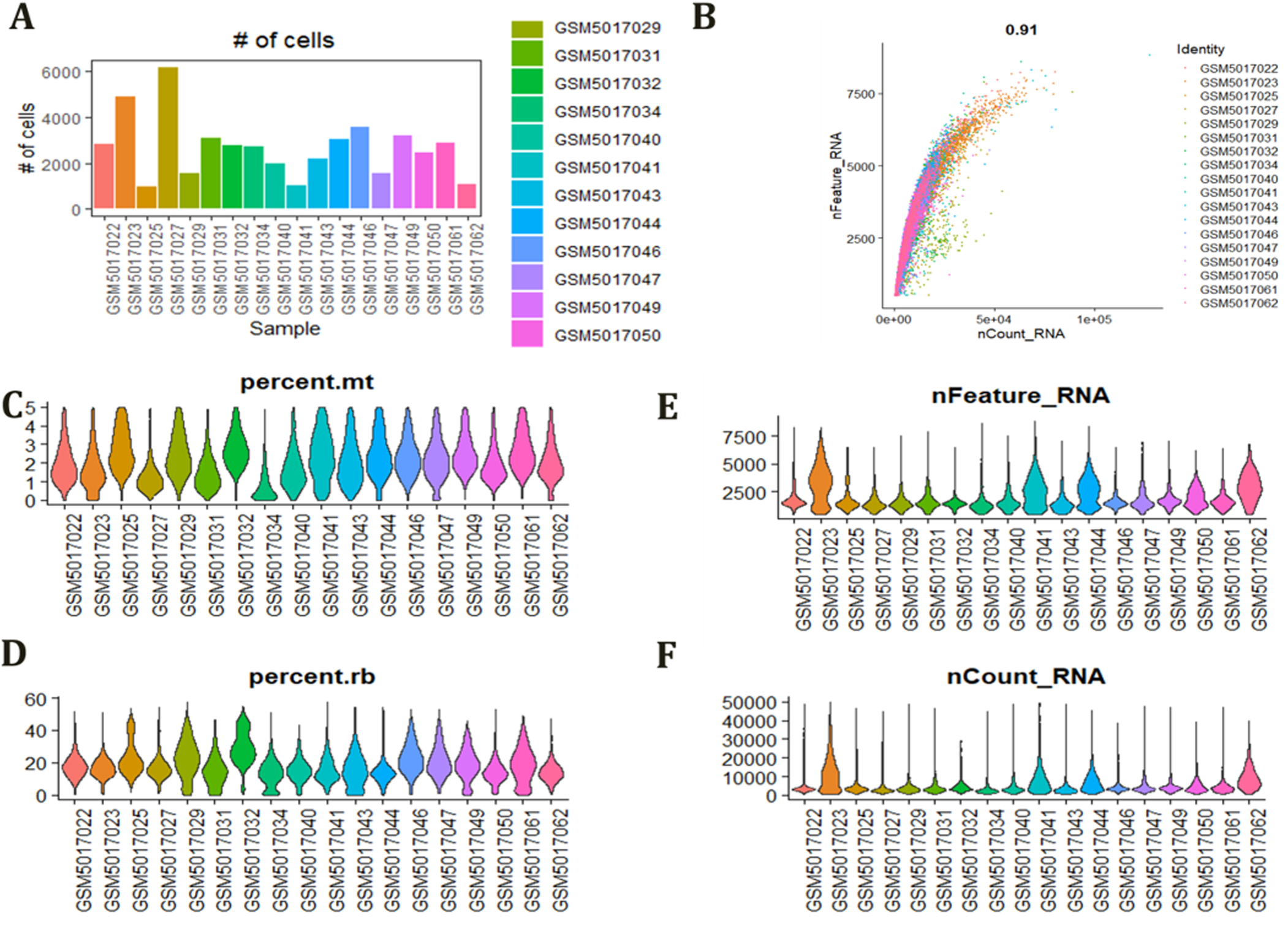
Clinical characteristics and quality control of the single-cell RNA-seq dataset. **(A)** Number of cells retained after quality control for each HPV-negative oral squamous cell carcinoma (OSCC) sample. **(B)** Relationship between the number of detected genes (nFeature_RNA) and unique molecular identifier (UMI) counts (nCount_RNA) across individual cells from each sample. **(C)** Distribution of mitochondrial transcript percentages (percent.mt) across samples. **(D)** Distribution of ribosomal transcript percentages (percent.rb) across samples. **(E)** Distribution of the number of detected genes (nFeature_RNA) across samples. **(F)** Distribution of unique molecular identifier (UMI) counts (nCount_RNA) across samples. Together, these quality control metrics demonstrate consistent sequencing quality and cellular complexity across the analyzed samples.

